# Dirac mixture distributions for the approximation of mixed effects models^⋆^

**DOI:** 10.1101/703850

**Authors:** Dantong Wang, Paul Stapor, Jan Hasenauer

## Abstract

Mixed effect modeling is widely used to study cell-to-cell and patient-to-patient variability. The population statistics of mixed effect models is usually approximated using Dirac mixture distributions obtained using Monte-Carlo, quasi Monte-Carlo, and sigma point methods. Here, we propose the use of a method based on the Cramér-von Mises Distance, which has been introduced in the context of filtering. We assess the accuracy of the different methods using several problems and provide the first scalability study for the Cramér-von Mises Distance method. Our results indicate that for a given number of points, the method based on the modified Cramér-von Mises Distance method tends to achieve a better approximation accuracy than Monte-Carlo and quasi Monte-Carlo methods. In contrast to sigma-point methods, the method based on the modified Cramér-von Mises Distance allows for a flexible number of points and a more accurate approximation for nonlinear problems.

## 1. INTRODUCTION

Cell-to-cell and patient-to-patient variability are ubiquitous and highly relevant phenomena. Cell-to-cell variability has a pronounced effect on the response to perturbations (Altschuler and Wu, 2010) and is considered as one of the major challenges in cancer therapy (Dagogo-Jack and Shaw, 2018). Patient-to-patient variability influences the processing of drugs and the response to treatments (Willmann et al., 2007).

Nonlinear mixed effects models (MEMs) are widely used to capture cell-to-cell (Karlsson et al., 2015; Llamosi et al., 2016; Fröhlich et al., 2018) and patient-to-patient variability (Tornoe et al., 2004; Willmann et al., 2007; Bastogne et al., 2009). These models provide a hierarchical description of populations (Pinheiro, 1994). Variability is encoded in parameter values, which are composed of fixed effects and random effects. The fixed effects are the same for the whole population, while the random effects vary between individuals. The biological process, e.g. cell signaling or pharmacokinetics, is encoded by a nonlinear function describing the dependence of outputs on parameters.

To simulate MEMs, the parameters of individual cells / patients are sampled and the nonlinear function is evaluated for these sampled parameters. The sampling essential yields a Dirac mixture distribution, which is simply a collection of Dirac components which can be weighted. To obtain robust estimates for the population statistics, e.g. mean and standard deviation, a substantial number of Monte-Carlo (MC) samples is required, causing a significant computational demand. To reduce the required number of samples, alternative Dirac mixture distributions can be employed. One alternative is the used of quasi Monte-Carlo (QMC) methods. However, to the best of our knowledge, this has not been published. In contrast to standard MC methods, QMC methods such as Sobol (Sobol, 1967) and Halton (Sobol, 1964) are based on low discrepancy sequences (Niederreiter, 1978). The use of low discrepancy sequences reduces the randomness and improves the robustness and convergence order.

Sigma point (SP) methods are an alternative to MC methods (Julier et al., 1995; Julier and Uhlmann, 2004; Menegaz et al., 2011; Lerner, 2002; van der Merwe, 2004; Charalampidis and Papavassilopoulos, 2011). These methods aim to approximate mean and covariance matrix of the parameters by sampling deterministically (van der Merwe, 2004). Then the approximated mean and covariance matrix of outputs are computed by propagating the samples through the model(Filippi et al., 2016; Loos et al., 2018). A problem is that the error is difficult to control for nonlinear models. Besides, for most SP methods, the number of samples is fixed or follows a certain formula, which makes it hard to flexibly improve the accuracy.

In this study, we propose the approximation of population statistics with Dirac mixture distributions constructed by minimizing a modified Cramér-von Mises Distance (CMD). The use of the CMD has been proposed in the context of state estimation, also known as filtering (Gilitschenski and Hanebeck, 2013). Here, we provide a first scalability analysis for CMD methods as well as a comparison with MC, QMC and SP methods for the analysis of nonlinear MEMs.

## 2. METHODS

### 2.1 Mixed effect models

We consider nonlinear MEMs as defined by Lindstrom and Bates (1990). The mixed effects parameter vector *ϕ* is a linear combination of fixed effects *β* and random effects *b*,

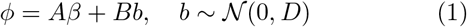

with design matrices *A* and *B*, and covariance matrix *D*. The response vector *y* is given by

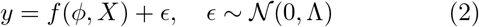

with a usually nonlinear function *f*, depending on the mixed effects parameter vector *ϕ* and the matrix of covariates *X*. Measurement noise is denoted by *ϵ* and follows a multi-variate normal distribution with covariance matrix Λ.

The nonlinear function *f* can have different forms, depending on the application. In systems and computational biology as well as pharmacology, *f* is frequently the solution operator to ordinary differential equations (ODEs) projected to a set of observables (Karlsson et al., 2015; Llamosi et al., 2016; Fröhlich et al., 2018). The ODEs often describe the dynamics of biochemical reaction networks with time-dependent concentration *c*, stoichiometric matrix *S* and reaction fluxes *v*(*c, ϕ*),

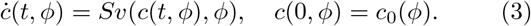

Measurements provide information about properties of this network,

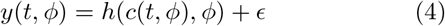

with *h* denoting the observation function which depends on the experimental device. In the case of a scalar observable *y* observed at time points *t_k_, k* = 1,…, *K*, this yields the nonlinear function

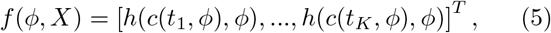

where the *h* depends on the solution of the ODE (3).

### 2.2 Evaluation of statistical moments of response vectors

We considered Dirac mixture distributions to evaluate the statistical moments of the response vector, in particular its mean 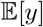 and covariance 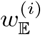. For MEMs, these methods use a set of random effects *b*^(*i*)^, *i* = 1,…,*n*, for which the corresponding mixed effects *ϕ*^(*i*)^ and responses *y*^(*i*)^ are evaluated. The estimators of the mean and covariance are

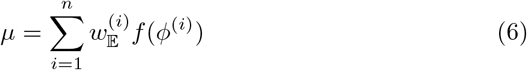

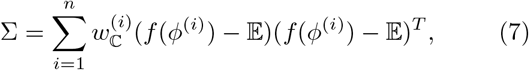

with weights 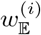 and 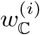. In the following, we outline the considered methods.

#### Remark

As measurement noise has no influence on the mean 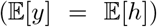 and an additive influence on the covariance 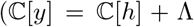, when the weights sum up to 1), it can be handled analytically. For the purpose of this study, we set it to zero (Λ = 0) to avoid that approximation errors are hidden.

**Monte Carlo (MC) methods** are the most commonly used approaches to determine integrals over probability distributions, such as statistical moments. The random effects *b*^(*i*)^ can be obtained by randomly sampling a multi-variate normal distribution 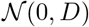. The weights are 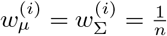.

MC methods yield unbiased results. However, this comes at the cost of reduced computational efficiency. It is well known that estimates obtained using small sample sizes *n* possess a large variance and that the convergence rate is only 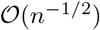. If the number of samples *n* is limited, e.g. due to the computational complexity of model simulation, even biased approaches might be beneficial.

**Quasi Monte Carlo (QMC) methods** address problems of MC methods by using low discrepancy sequences. These sequences are more regular than random samples used by MC methods (Fig. 1a). This improves the convergence rate to 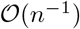 (Caflisch, 1998). The points of the sequence are used in the same way as MC samples.

**Fig. 1.**
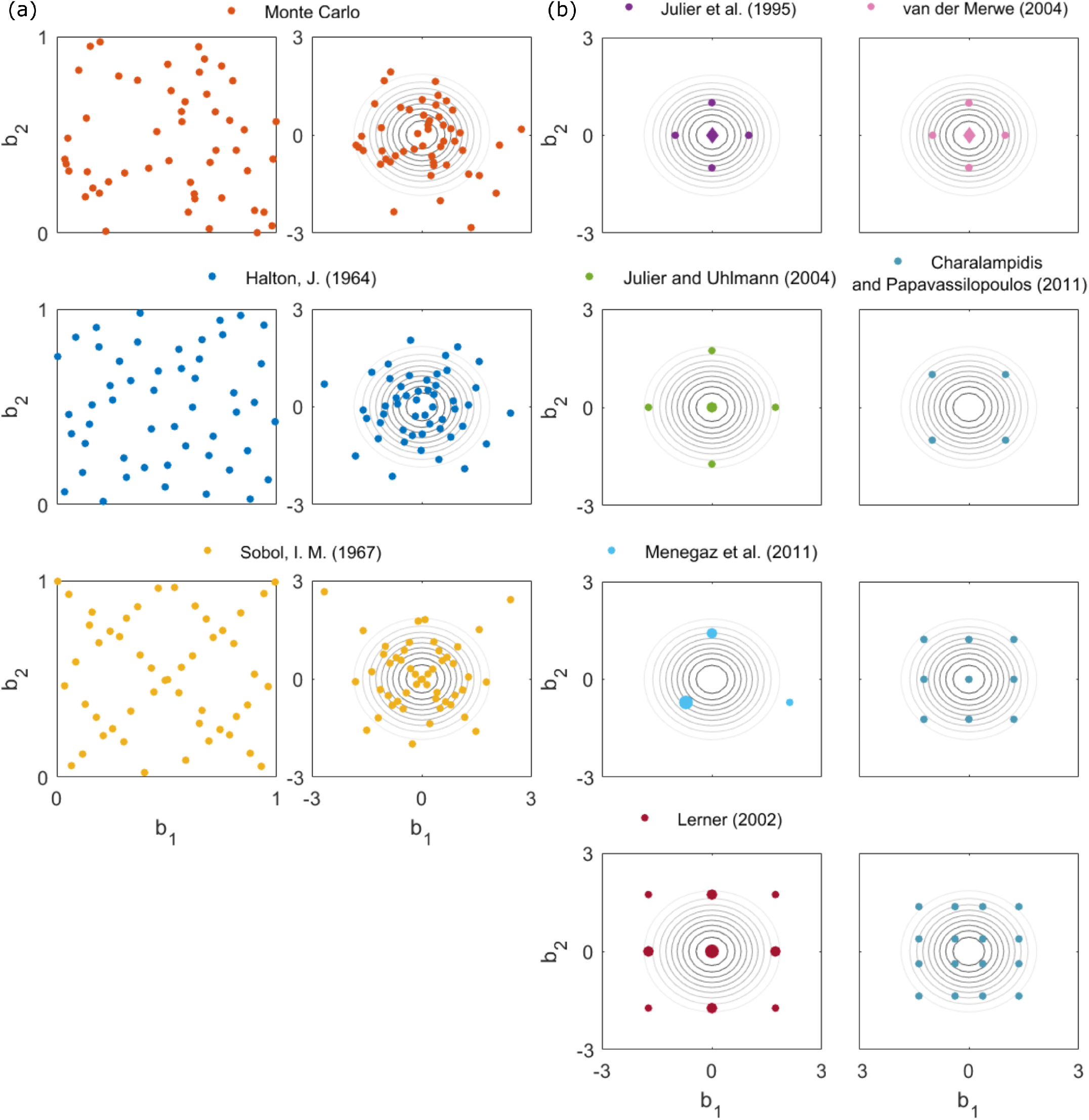
Illustration of MC, QMC and SP methods in two dimensions. (a) Random sequence used by the MC method, and Halton and Sobol sequence used by QMC methods: (left) sequences in the unit cube, and (right) transformed sequences to approximate standard normal distribution. As a reference, level sets of the standard normal distribution are shown as gray contour lines. (b) Point distributions and weights for different SP methods. Positive point weights are indicated by a circle, negative weights by a diamond. The absolute values of the weights is encoded by the marker size (small markers for small absolute values, and large markers for large absolute values). Among others, it is possible for Charalampidis method to have different sample sizes: *n* = *θ^L^*, where *θ* can be any integer and *L* is the dimension.

In this study, we consider Halton and Sobol sequences. As these sequences are defined on the unit cube, we transformed them to representative sequences for the standard multi-variate normal distribution using the corresponding cumulative distribution function. The application of the Mahalanobis transform – using the matrix square-root of *D* – to these sequences yields samples for the random effects *b*^(*i*)^ with the desired covariance. The Halton and Sobol sequences performed similar (Fig. 2), therefore we only show results for Halton sequences.

**Fig. 2.**
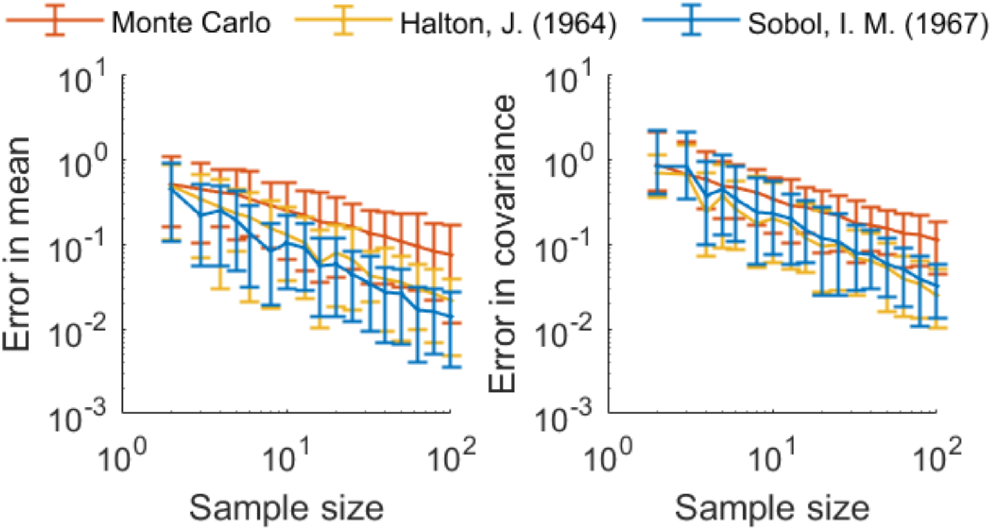
Absolute difference between true moments (mean and covariance) and sample moments. The results of MC and QMC methods for the standard normal distribution are depicted. The interval between the 5th and the 95th percentile of the errors across 1000 realizations are depicted.

**Sigma point (SP) methods** aim to improve the approximation accuracy achieved for small sample sizes. Therefore, the sequences are designed such that they yield the exact statistical moments of the normal distribution up to a particular order. This ensures good approximation properties if *f* is close to linear in the region of interest.

In contrast to MC and QMC methods, SP methods use non-uniform weights 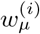 and 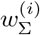. The weights of mean and covariance might differ 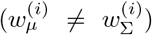 and can be negative for some points. Furthermore, the number of points *n* is usually rather small, ensuring computational efficiency. However, a continuous refinement of the SP approximation is not possible as only certain numbers of sample points are feasible.

In this study, we consider six different SP methods (Julier et al., 1995; Julier and Uhlmann, 2004; Menegaz et al., 2011; Lerner, 2002; van der Merwe, 2004; Charalampidis and Papavassilopoulos, 2011). These are in our opinion the most widely used SP methods. The methods differ in the number of points n, or their locations and/or weights (Fig. 1b). For details on the selection of the point locations, we refer to the original publications. An overview over the SP methods is provided in Table 1.

**Table 1.** Properties of SP and CMD methods. The references, number of points *n* (for an *L*-dimension random effect vector) and moments of Gaussian distribution which are matched precisely are listed.

| Method | Number of points | Exact moments |
| --- | --- | --- |
| Julier et al. (1995) | $2L + 1$ | 1st, 2nd |
| Julier and Uhlmann (2004) | $2L + 1$ | 1st, 2nd |
| Menegaz et al. (2011) | $L + 1$ | 1st, 2nd |
| Lerner (2002) | $2L^2 + 1$ | 1st, 2nd |
| van der Merwe (2004) | $2L + 1$ | 1st, 2nd |
| Charalampidis and Papavassilopoulos (2011) | $\theta^L$ | 1st, 2nd |
| CMD | flexible | 1st |

In this study, we propose **CMD methods** for the approximation of statistical moments of nonlinear MEMs. This method constructs a representative set of points by minimizing the modified CMD,

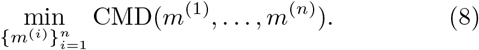

The modified CMD has been introduced by Gilitschenski and Hanebeck (2013) and is based on a local cumulative distribution function (Hanebeck and Klumpp, 2008). It measures the discrepancy between a Dirac mixture distribution and a multi-variate standard normal distribution. To samples with a given covariance matrix, the Mahalanobis transform is applied to the solution of (8) to obtain *b*^(*i*)^, *i* = 1,…, *n*.

To the best of our knowledge, CMD methods are not widely used and have not been applied in MEMs. Yet, CMD methods possess several advantageous properties: (1) CMD methods allow – similar to MC and QMC methods – for any discrete number of samples (Fig. 3a), which enables a control of approximation accuracy; and (2) they provide robust approximations at low sample sizes.

**Fig. 3.**
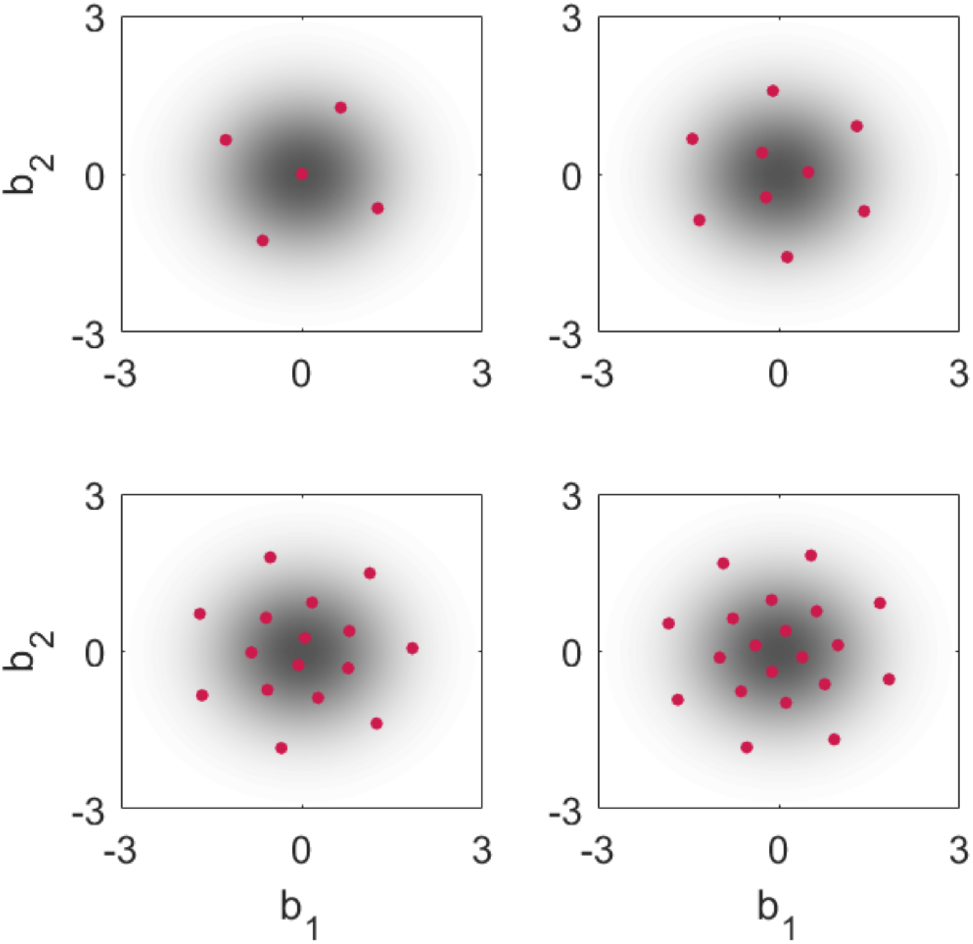
**Location of point sets obtained using the CMD method** with the sample size *n* = 5, *n* = 9, *n* = 16, *n* = 20.

CMD methods consider the distribution and not particular moments. Accordingly, even for linear models the mean might be off. To address this problem, we enforced a zero mean by estimating only *n* − 1 points and setting 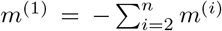. Furthermore, we generalized CMD methods by Gilitschenski and Hanebeck (2013) to allow for non-uniform weights. Due to page limitations, the derivation of the generalized CMD method is provided in the open-source MATLAB toolbox SPToolbox: https://github.com/ICB-DCM/SPToolbox.

### 2.3 Implementation

We implemented the MC, QMC, SP and CMD methods in the SPToolbox. For the construction of Sobol and Halton sequences, the MATLAB built-in functions `sobolset` and `haltonset` are used.

To solve optimization problem (8), we employ multi-start local optimization implemented in the open-source MATLAB toolbox PESTO (Stapor et al., 2018). The local optimization is performed using the MATLAB function `fmincon`. This optimizer is provided with analytical gradients obtained by differentiating the objective function with respect to the optimization variables. The starting points for the optimization are obtained using latin hypercube sampling. The sampling and the optimization used for the locations with lower bounds = −3 and upper bound = 3, as this regime contains for standard normal distribution more than 99% of the probability mass.

## 3. RESULTS

We provide an assessment of the different methods using two test problems:

- Quadratic function (M1): 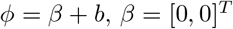 for *i* = 1, 2 with *ϕ* = *β* + *b, β* = [0, 0]^*T*^ and 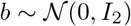
- Hill-type function (M2): 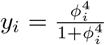 for *i* = 1, 2 with *ϕ* = *β* + *b, β* = [0, 0]^*T*^ and 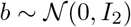
and two application problems. The first application problem is a model of a conversion reaction model (M3):
- Conversion reaction model (M3): *y* = *x*_2_

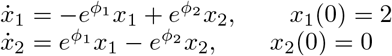

with *ϕ* = *β* + *b, β* = [0, 0.3]^*T*^ and 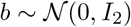

Conversion reactions are common motives in biological processes. The second application example is a model for JAK-STAT signaling (M4) based on Swameye et al. (2003). This model is widely used for method evaluation (Raue et al., 2009; Hass et al., 2016; Fröhlich et al., 2016) and possesses 9 state variables, 16 parameters and 3 observables. We assume that 5 parameters differ between cells.

The implementation of all models is provided in the SPToolbox.

To compute the ground truth of the statistical moments for the test and application problems, we used MC sampling with 100,000 points. Since outputs for the application examples are multi-dimensional, average values of absolute differences to true mean and true variance were computed.

### 3.1 Evaluation and extension of the CMD method

As the calculation of random effects in the CMD method requires the solution of an optimization problem, the scalability of the approach is unclear. To assess it, we evaluate the average computation time per local optimization (Villaverde et al., 2018), which is the overall computation time divided by the number of local optimization runs achieving the best objective function value. We found that the computation time scales roughly linearly with the number of points n (Fig. 4a) and linearly with the dimension of the random effect vector (Fig. 4b). Accordingly, the computation time required for optimization increases substantially with both problem dimensions.

**Fig. 4.**
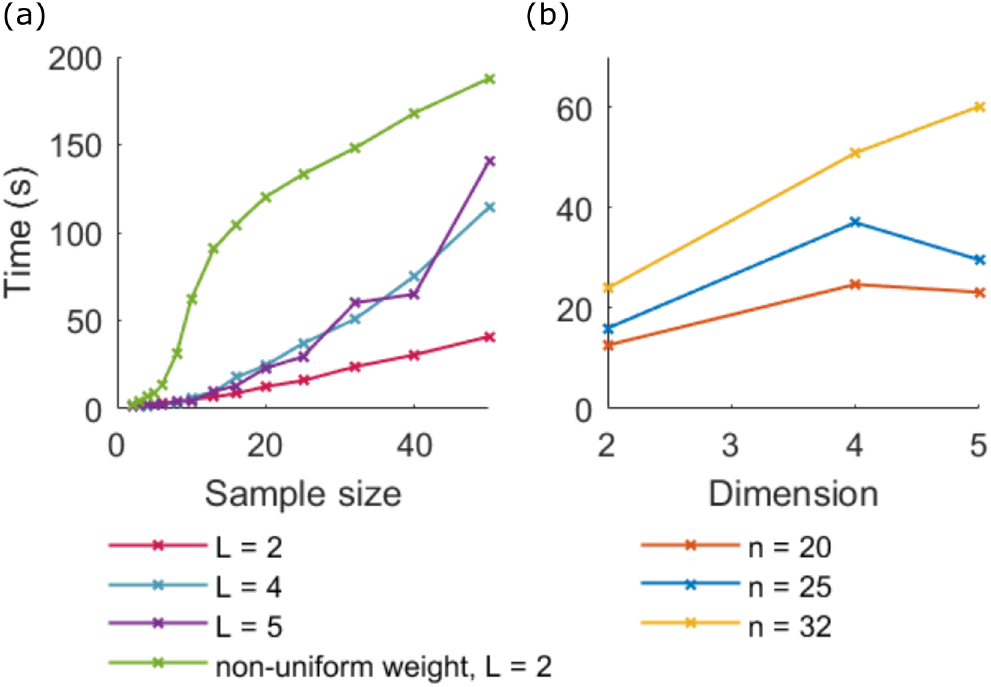
Scalability of the CMD method. Average computation time per converged start for (a) different number of points and (b) different dimensions is depicted.

Non-uniform weighting of the points was considered to reduce the number of points. Therefore, we optimized points and weights together, using analytically derived expressions for the gradients. We found that point locations change (see Fig. 3 vs. 5) and that points close to the center of the distributions tend to have higher weights (Fig. 5). The value of the modified CMD improved marginally by introducing the weights (for *n* = 8: uniform CMD = 0.0261 and non-uniform CMD = 0.0236). This improvement comes at a substantially increased computation time (Fig. 4a).

**Fig. 5.**
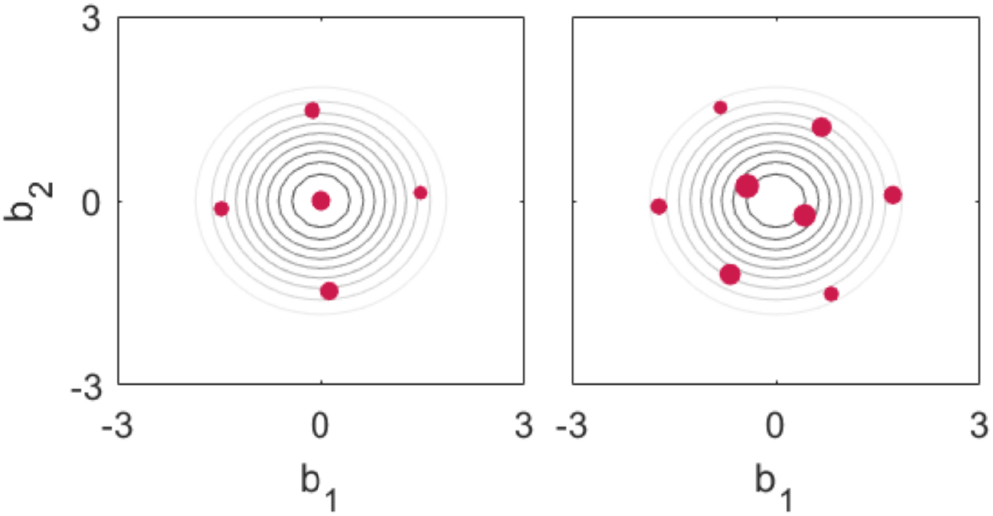
**Location of weighted CMD points** with sample size *n* = 5 and *n* = 8. The values of the weights are encoded by the marker size.

An analysis of the optimization problem (8) reveals that the optimal point is not unique. Optimized point properties can be swapped and the point sets can be rotated around the origin without changing the objective function values. While points swaps can not have any influence on the approximation of the moments, the rotation angle can have an influence when *f* is nonlinear. However, our analysis shows that the standard deviation of the approximation error was smaller than its mean (Fig. 6a,b), suggesting that the effect is not too important. As the point sets are structured, the approximation error shows a periodic behavior. The period length depends on the number of points (Fig. 6a).

**Fig. 6.**
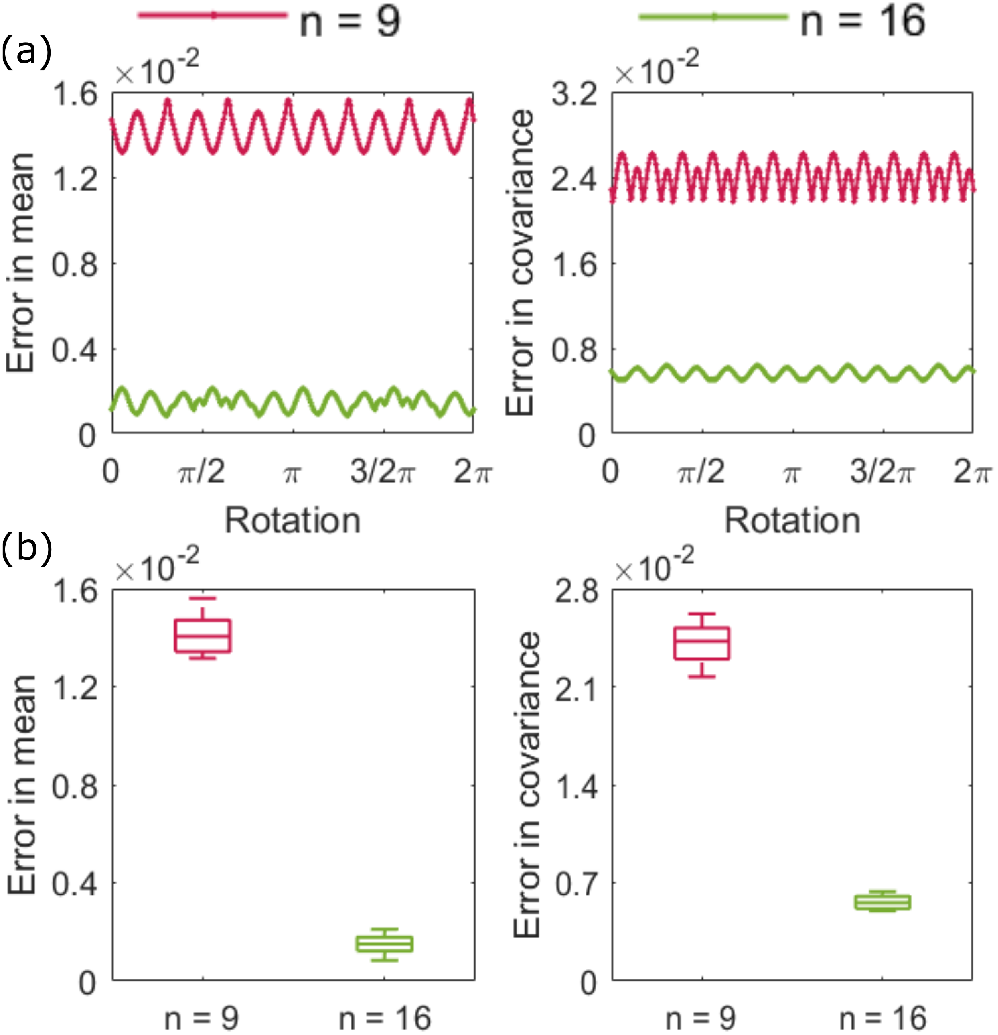
Influence of rotation angle for problem M2. (a) Absolute difference between true mean and variance and the approximations obtained using the CMD method with rotated point sets. (b) Box plot showing the variability induced by the rotation.

### 3.2 Approximation properties of Dirac mixture distributions

To assess the approximation accuracy of the MC, QMC, SP and CMD methods, we applied them to the test and application problems. For the MC, QMC and CMD methods, as well as the SP method by Charalampidis and Papavassilopoulos (2011), multiple point numbers were considered to study the convergence rates. For all problems we determined the absolute differences of mean and variance compared to the ground truth.

For M1, the mean computed by the SP methods is exact (by construction). For all models, the CMD method with uniform weights achieves a better accuracy than the Halton QMC method and SP methods (Fig. 7). In addition, compared to the Halton method, CMD methods show (1) similar or better convergence rates and (2) smaller confidence intervals. The latter implies an improved stability of the approximation and an independence from the rotation angle. CMD methods with uniform and nonuniform weighting achieve similar accuracy.

**Fig. 7.**
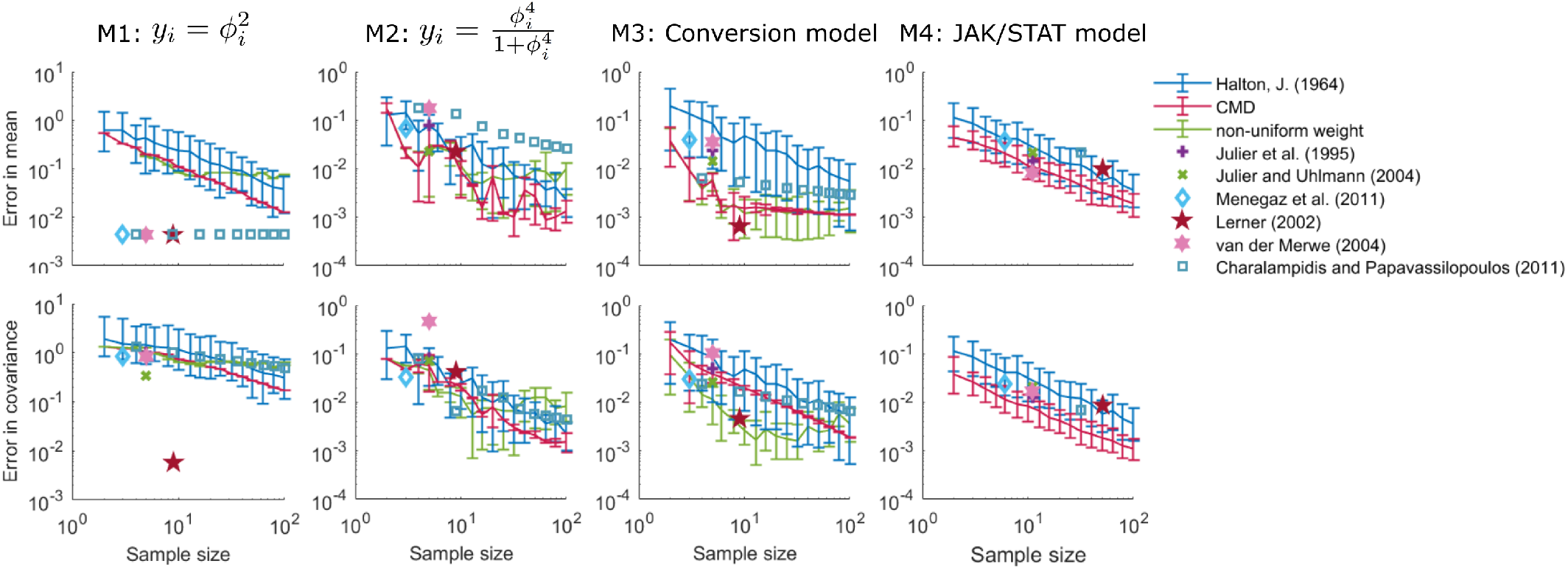
Assessment of the approximation accuracy for test and application problems. The absolute difference between approximated moments and ground truth is depicted: (upper) mean and (lower) covariance.

## 4. DISCUSSION

In this study, we perform an evaluation of MC, QMC, SP and CMD methods. For CMD methods – which are not commonly applied for MEMs – we perform a first scalability study and assess the influence of non-uniform weights and rotation angles.

Our results show that compared to the SP methods, CMD methods are more flexible since a continuous refinement is possible, and possess a better accuracy when the model is highly nonlinear. Compared to MC and QMC methods, CMD methods have better convergence and lower variance, especially for more complicated models. Nonuniform weighting does not seem to provide a substantial benefit. Overall, our results demonstrate the potential use of CMD methods.

